# A commonly used photosynthetic inhibitor fails to block electron flow to photosystem I in intact systems

**DOI:** 10.1101/857177

**Authors:** Duncan Fitzpatrick, Eva-Mari Aro, Arjun Tiwari

## Abstract

In plant science, 2,4-dinitrophenylether of iodonitrothymol (DNP-INT) is frequently used as an alternative to 2,5-dibromo-6-isopropyl-3-methyl-1,4-benzoquinone (DBMIB) to examine the capacity of plastoquinol and semiquinone to reduce O_2_. DNP-INT is considered an effective inhibitor of the photosynthetic electron transfer chain (PETC) through its binding at the Q_0_ site of Cyt-*b6f*. The binding and action of DNP-INT has been previously characterized spectroscopically in purified Cyt-*b6f* complex reconstituted with Plastocyanin, PSII membranes and plastoquinone, as well as in isolated thylakoids based on its property to block MV-mediated O_2_ consumption. Contrary to the conclusions made from these experiments, we observed clear reduction of P700^+^ in samples incubated with DNP-INT during our recent investigation into the sites of oxygen consumption in isolated thylakoids. Therefore, we carried out an extensive investigation of DNP-INT’s chemical efficacy in isolated thylakoids and intact leaves. This included examination of its capacity to block the PETC before PSI, and therefore its inhibition of CO_2_ fixation. P700 redox kinetics were measured using Dual-PAM whilst Membrane Inlet Mass Spectrometry (MIMS) was used for simultaneous determination of the rates of O_2_ evolution and O_2_ consumption in isolated thylakoids and CO_2_ fixation in intact leaves, using two stable isotopes of oxygen (^16^O_2_,^18^O_2_) and CO_2_ (^12^C,^13^C), respectively. Based on these investigations we confirmed that DNP-INT is unable to completely block the PETC and CO_2_ fixation, therefore its use may produce artefacts if applied to isolated thylakoids or intact cells, especially when determining the locations of reactive oxygen species formation in the photosynthetic apparatus.

## 1. Introduction

Life in all three domains is sustained by membrane protein complexes participating in circuits that couple redox reactions and proton pumping, with the generation of ATP and NAD(P)H. The strategic application of chemical inhibitors, in isolation and in concert, has proven invaluable to study the roles, components and mechanisms of electron transport chains. In studies of photosynthetic systems, 3-(3,4-dichlorophenyl)-1,1-dimethylurea (DCMU), inhibitor of plastoquinone (PQ) reduction by PSII (Trebst 2007) (see Figure 1) and 2,5-Dibromo-6-isopropyl-3-methyl-1,4-benzoquinone (DBMIB), inhibitor of PQ-oxidation by cytochrome (Cyt)-*b6f* complex (Trebst 2007) (see Figure 1) have been the most regularly used and commercially available inhibitors. A drawback of the quinol base of DBMIB is that it possesses intrinsic redox functionality that in excess concentrations provides endogenous electron transport capacity able to bypass Cyt-*b6f* (Chain and Malkin 1979). To counter this, 2,4-dinitrophenylether of iodonitrothymol (DNP-INT) was synthesized as a redox inert surrogate with characteristics otherwise similar to DBMIB (Trebst and others 1978).

**Figure 1.**
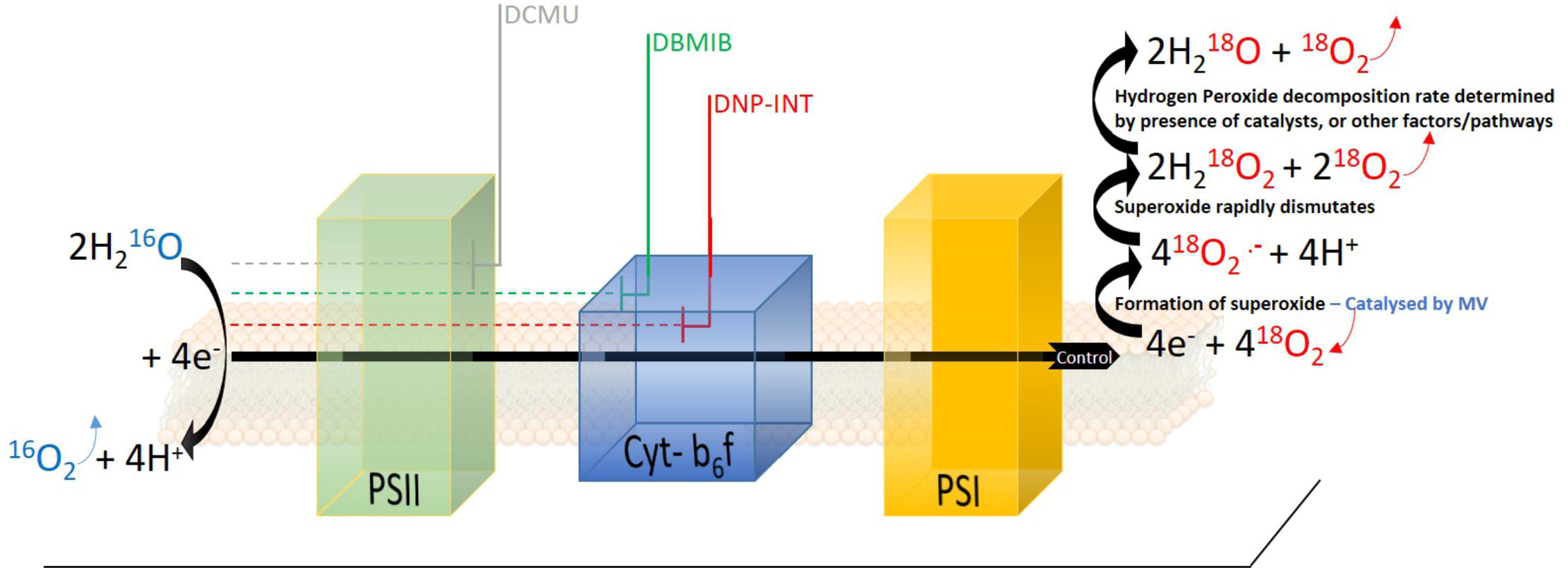
Schematic diagram of photosynthetic electron transport chain in isolated thylakoid samples when natural ^16^O_2_ isotope (Blue) was diminished significantly from the measuring chamber and enriched with the heavy ^18^O_2_ isotope (Red). One molecule of ^16^O_2_ (Blue) and four electrons are produced through oxidation of two water molecules at PSII. Electrons are transferred to four molecules of ^18^O_2_ at the acceptor side of PSI (potentially also within PQ pool), forming four superoxide molecules (step catalyzed by Methyl Viologen, MV). The superoxide dismutates rapidly into two hydrogen peroxide molecules, releasing two molecules of ^18^O_2_. Further decomposition of H_2_O_2_ to H_2_O and O_2_ is the final step in the electron transfer pathway of the water-water cycle. Approximate locations of inhibition of electron transport targeted by DCMU, DBMIB and DNP-INT are indicated in the figure.

Significant effort has been committed to characterize the inhibitory action of DNP-INT in the photosynthetic electron transfer chain (PETC). Absorption spectroscopy measurements of purified Cyt-*b6f* complex reconstituted with plastocyanine (PC), PQ, photosystem (PS)I and PSII in the reaction medium (Lam and Malkin 1983; O’Keefe 1983), along with low temperature electron paramagnetic resonance (EPR) spectroscopy of purified Cyt-*b6f* complex (Malkin 1986; Roberts and Kramer 2001) have shown that DNP-INT blocks oxidation of plastoquinol and reduction of PC by binding at the Q_0_ site of *Cyt*-*b6f*. Furthermore, functional measurements have been used to demonstrate the effectiveness of DNP-INT in blocking linear electron transfer in more complex samples, such as isolated thylakoids. To this end, the oxygen (O_2_) consumption rate upon exposure of thylakoid samples to light was measured with a Clarke type O_2_ electrode in the absence of artificial electron acceptors (Khorobrykh and Ivanov 2002). These experiments demonstrated similar O_2_ uptake rates between untreated and DNP-INT incubated thylakoid samples, yet the addition of Methyl Viologen (MV), a strong catalyst of O_2_ reduction at PSI acceptor side (see Figure 1), increased Net O_2_ uptake rates only in samples lacking DNP-INT. This was interpreted as strong evidence that DNP-INT blocks electron flow via PSI to MV, and therefore the chemical effectively truncates PSI from the PETC. Furthermore, similarity in O_2_ uptake rates observed between untreated and DNP-INT treated samples suggested the presence of superoxide forming pathways, similar to that in PSI, also associated with the reduced PQ pool and *cyt*-*b6f* complex (Khorobrykh and Ivanov 2002; Mubarakshina and Ivanov 2010; Borisova-Mubarakshina, Naydov and Ivanov 2018).

The results from O_2_ electrode measurements with isolated thylakoids were interpreted as the ability of DNP-INT to allow semiquinone formation, whilst simultaneously blocking cyt-*f* reduction (Mubarakshina and Ivanov 2010), which made DNP-INT a critical tool in truncating the PETC at Cyt-*b6f*. It has resulted in the wide use of DNP-INT as an alternative to DBMIB for truncation of PSI from the PETC, particularly in works to explore reactive oxygen species (ROS) formation and scavenging pathways involving both reduced PQ-pool and semiquinones (Khorobrykh and Ivanov 2002; Mubarakshina and Ivanov 2010; Borisova-Mubarakshina, Naydov and Ivanov 2018; Heyno and others 2009). Based on these characterizations, use of DNP-INT has provided a range of significant findings in plant (Stepien and Johnson 2009) and algal physiology (Barbagallo, Finazzi and Forti 1999), bioenergetics (Malnoe and others 2011), bio-fuel applications (Mus and others 2005), biochemical characterizations of ETC components (Krieger-Liszkay, Kienzler and Johnson 2000) and efforts to determine sites and activity of ROS formation/quenching (Khorobrykh and Ivanov 2002; Mubarakshina and Ivanov 2010; Borisova-Mubarakshina, Naydov and Ivanov 2018; Heyno and others 2009; Vetoshkina and others 2017).

In our current work, we attempted to determine the contribution of PSI in overall light-induced O_2_ reduction from isolated thylakoids and intact leaf discs. For this, we planned to truncate the ETC with inhibitors such as DBMIB and DNP-INT. As a control for successful and complete truncation of PSI from the PETC under our specific experimental conditions, we measured both isolated thylakoid and intact leaf samples with a Dual PAM to record the redox kinetics of P700. However, DNP-INT failed to block linear electron transport to PSI in both sample types. After controlling for the chemical structure of our commercially procured DNP-INT with ^1^HNMR spectroscopy, and following our successful reproduction of the frequently published O_2_ electrode results (Khorobrykh and Ivanov 2002; Vetoshkina and others 2017) described above, we embarked on a thorough investigation of PSI redox kinetics and photosynthetic gas exchange in DNP-INT treated samples. To confirm whether DNP-INT truly truncates PSI from the PETC, we compared the redox kinetics of P700 measurements incubated with DNP-INT and other well characterized PETC inhibitors. Membrane Inlet Mass Spectrometry (MIMS) was used to distinguish the O_2_ consuming and O_2_ producing reactions occurring simultaneously in isolated thylakoid samples to test conclusions drawn from O_2_ electrode data, and was used then to measure rates of CO_2_ fixation in leaf discs infiltrated with DNP-INT.

## 2. Materials and Methods

### 2.1 Leaf samples and isolation of thylakoids

Thylakoids were isolated from 6 week old *Arabidopsis thaliana* plants, grown at 16/8 h dark/light cycle at 120 μmol photons m^−2^ s^−1^ at atmospheric CO_2_. Thylakoids were isolated as described earlier (Tiwari and others 2016) except all measurements were performed with freshly isolated thylakoids in buffer containing: 330 mM Sorbitol, 5 mM MgCl_2_, 10 mM NaCl, 5 mM NH_4_Cl, 50 mM Hepes pH 7.6. Leaf measurements in MIMS and Dual-PAM were taken from the same plants.

### 2.2 MIMS measurements of isolated thylakoid membranes

Freshly isolated thylakoid samples of known chlorophyll concentration were stored on ice in darkness. A Sentinel-PRO magnetic sector mass spectrometer (Thermo-Fisher USA) was employed to collect masses 32 and 36 with a total cycle time of approximately 4.5 seconds. For each run sufficient measurement buffer (containing: 330 mM Sorbitol, 5 mM MgCl_2_, 10 mM NaCl, 5 mM NH_4_Cl, 50 mM Hepes pH 7.6) was loaded into the cuvette equilibrated and calibrated to 25 °C, thylakoids were added (equivalent to approximately 50 μg chlorophyll) to a final volume of 1000 μl. In darkness the sample was briefly purged with N_2_ to reduce the background ^16^O_2_ signal before a bubble of ^18^O_2_ (Cambridge Isotope Laboratories Inc, UK) was loaded into the stirring liquid, bringing the concentration of the heavier isotope up to approximately 150 nmol ml^−1^. The bubble was removed and inhibitors were injected at this moment (either 10 μM DCMU, 10 μM MV, 10 μM DBMIB, 10 μM DNP-INT) and samples equilibrated in darkness for five minutes before data acquisition was started. Samples were illuminated via halogen lamp (Dolan Jenner, USA) at 120 μmol photons m^−2^ s^−1^. At the end of each run, the Chl concentration of the sample was determined in triplicate using the Porra Method (Porra, Thompson and Kriedemann 1989) in 90% MeOH to ensure accurate normalization of rates between samples. The cuvette was washed thoroughly with multiple rinses of 70% Ethanol followed by MQ H_2_O when changing between inhibitors to avoid cross contamination. This was checked by running controls. All data was analysed and fluxes calculated with equations described in Beckmann et al (Beckmann and others 2009).

### 2.3 MIMS measurements of leaf discs

Leaf discs (14 mm) were cut from detached leaves and floated in darkness in either H_2_O (control) or H_2_O + 10 μM DBMIB or H_2_O + 10 μM DNP-INT. Samples were incubated overnight in darkness at 25 °C. For measurements, all excess water from leaf surface was removed and a smaller 12.5 mm disc was cut from the larger disc to remove the old edge. This was loaded into an in-house built stainless steel cuvette of 1000 μl volume, equilibrated and calibrated at 25 °C, using Teflon (Hansatech) membrane to separate the sample space from the high vacuum line of the Mass Spectrometer. With the disc maintained in darkness, the cuvette was purged with N_2_ to remove atmospheric ^16^O_2_ and ^12^CO_2_ before ^18^O_2_ (as above) and ^13^CO_2_ (Sigma Aldrich, USA) were injected to approximately 3 % and 2 % by volume respectively. Discs were kept for approximately five minutes in darkness inside the cuvette to ensure isotopic equilibrium in the system before the measurement started. The instrument recorded m/z 32, 36, 44 and 46 with a time resolution of approximately 6.5 seconds. After four minutes dark a halogen light directed via a liquid light guide illuminated the samples at 120 then 520 μmol photons m^−2^ sec^−2^. All data was analysed and fluxes calculated with equations described in Beckmann et al (Beckmann and others 2009).

### 2.4 P700 redox kinetics measurements

P700 redox kinetics were measured with detached leaves and isolated thylakoids using Dual-PAM-100 or Dual-Klass NIR (Walz, Germany). Isolated thylakoids in Dual Pam were measured at 100 μg Chl ml^−1^ using liquid sample holder. The P700 was oxidised under continuous far red (FR) light. Activating two short pulses of saturating actinic light *i.e.* single turnover (ST) (50μs) and multiple turnover (MT) (50ms) pulses, over the FR light oxidised P700, induced partial P700 re-reduction by electrons from PSII (Tiwari and others 2016). Infiltration of inhibitors was conducted in the same manner as described for MIMS, however, the incubation time for leaves was only one hour.

### 2.5 O_2_ Electrode measurements

Thylakoid membranes prepared for MIMS measurements were also submitted to the Clark-type O_2_ electrode (Hansatech, UK) measurements. 15 μg Chlorophyll ml^−1^ of sample was injected to 1000 μl of the same measurement buffer used in MIMS measurements. Inhibitors were added as described in the text. Samples were loaded in darkness and after one minute of dark data acquisition the halogen lamp was turned on (800 μmol photons m^−2^ s^−1^) for four minutes (total five minutes).

## 3. Results and discussion

### 3.1 P700 redox kinetics suggested DNP-INT does not truncate PSI from the PETC

In attempts to differentiate rates of O_2_ reduction between the acceptor side of PSI and the reduced PQ pool/Cyt-*b6f* complex, we have tested the efficacy of DBMIB and DNP-INT, two well characterized chemicals used to truncate PSI from the PETC, to validate that PSI was not reduced during our measurements. To examine this question we used a Dual PAM to measure the redox kinetics of P700 from both intact leaf and isolated thylakoid samples incubated with these inhibitors, and further compared them with DCMU as an unambiguous control for the inhibition of all intersystem electron transport between PSII and PSI (Figure 2A,B).

**Figure 2.**
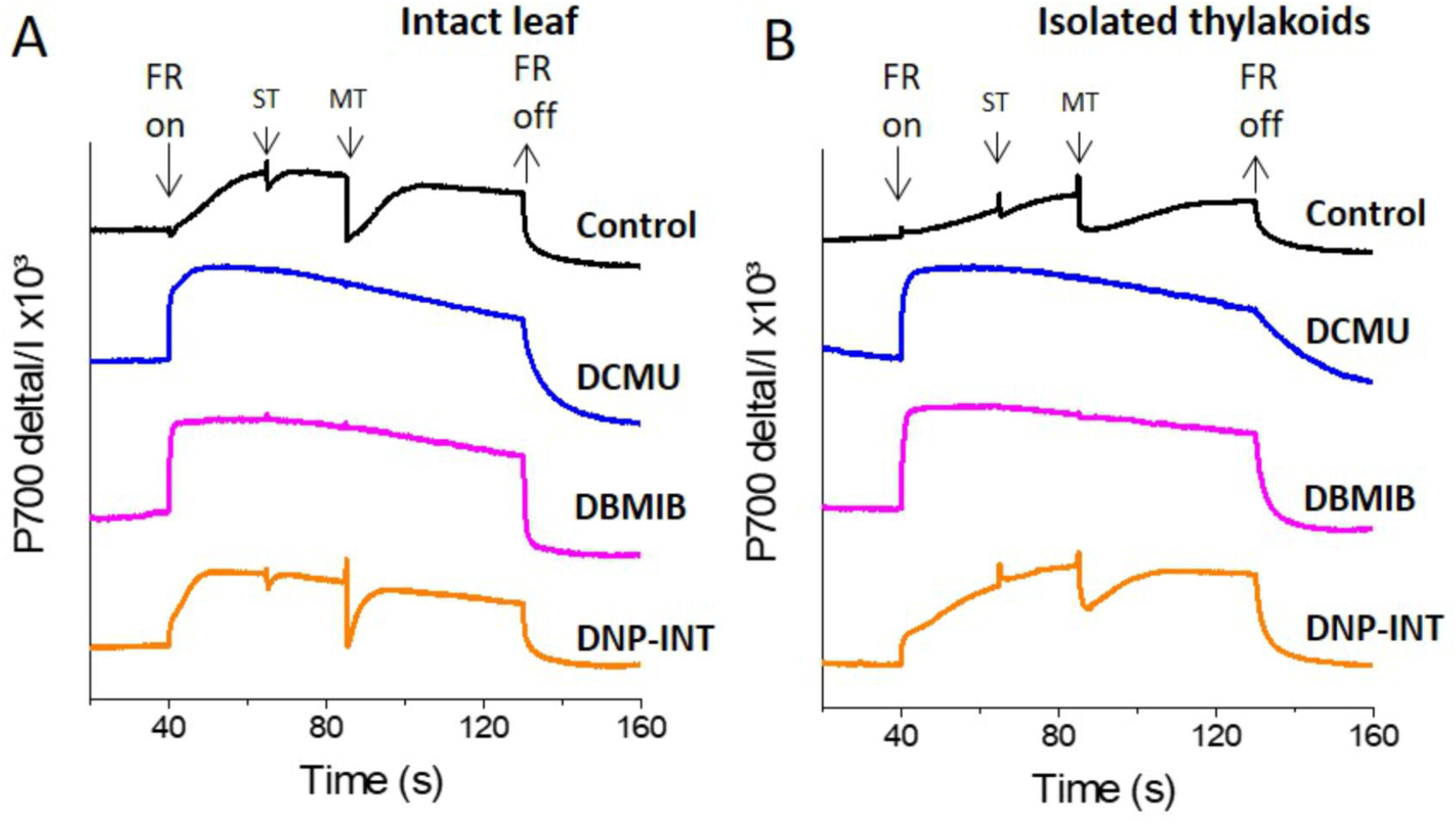
Comparison of P700 redox kinetics from DCMU, DBMIB and DNP-INT infiltrated intact leaf and isolated thylakoid samples, measured with Dual Pam. **(A)** P700 redox kinetics of intact leaves; **(B)** P700 redox kinetics of isolated thylakoids. Uncoupled thylakoid samples were measured in liquid sample holder cuvette of Dual Pam at a chl concentration of 100 μg Chl ml^−1^ in buffer containing only 330 mM Sorbitol, 5 mM MgCl2, 10 mM NaCl, 5 mM NH4Cl, 50 mM Hepes pH 7.6 or with each chemical modulator at 10 μM as indicated in figure. Leaves were infiltrated in darkness using water for the control sample and all inhibitors were used at 10 μM concentration. Representative curves are average of minimum n=5 samples. FR = P700 oxidising Far Red light. ST = Single turnover (50 μs) actinic light pulse, MT = multiple turnover (50 ms) saturating light pulse.

Following a dark period, addition of a constant Far Red (FR) light oxidized P700 to P700^+^ in all samples (Figure 2A,B). Transient re-reduction of P700^+^ to P700 by PSII derived electrons was achieved in inhibitor-free controls (Black lines, Figure 2A,B) via superimposition of saturating single turnover (ST) and multiple turnover (MT) actinic light pulses over the oxidising FR light. Such transient re-reduction requires, and therefore tested, the function of the entire intersystem ETC (Fan and others 2016; Tiwari and others 2016). Extinguishing the FR light allowed P700^+^ to re-reduce to P700, at varying rates, in all samples. In a clear demonstration of successful PSI truncation from the rest of the PETC, the level of P700^+^ was largely unaffected following the ST and MT actinic pulses in both leaf and thylakoid samples treated with 10 μM DCMU (Blue lines) and 10 μM DBMIB (Pink lines). Unexpectedly, leaf and thylakoid samples treated with 10 μM DNP-INT (Orange lines) behaved in a manner resembling untreated controls. Although smaller in amplitude as compared to the controls, these dips (re-reduction of P700^+^) produced by ST and MT actinic pulses in the P700^+^ curves of DNP-INT treated leaf and thylakoid samples were undeniably present.

This suggested that electrons liberated by PSII were able to transiently reduce P700^+^ to P700, whilst the smaller amplitude of the dips suggested that the number of electrons transferred to P700^+^ in DNP-INT treated samples was impaired, compared to untreated samples. We also used a Dual Klass NIR to measure the redox kinetics of PC (Figure S1 in Supplementary material), which accepts electrons exclusively from Cyt-*f*. For all treatments, the PC kinetics essentially matched those discussed for the P700 data. This suggests that use of DNP-INT at 10 μM in isolated thylakoid samples did not block reduction of Cyt-*f*, and therefore P700 was still actively coupled to PSII, as opposed to the clear inhibition of PC and P700^+^ re-reduction produced by the treatment of samples with DCMU and DBMIB.

### 3.2 Our DNP-INT stocks and isolated thylakoids reproduce published data

The concentration, quality and activity of our DNP-INT stocks were checked before making further investigations into the results of the Dual PAM data. Although the 10 μM DNP-INT concentration used in our protocol was at the high end of published protocols, and five times the “minimum concentration required for complete inhibition” of 2 μM (Trebst and others 1978), we tested the possibility that we had under-dosed the DNP-INT samples by conducting a concentration response curve (See Figure S1 in supplementary materials). In this experiment, DNP-INT failed to block the transient re-reduction of P700 caused by actinic light pulses in both the leaf and thylakoid samples from 2 μM to 500 μM concentration. That is 1000x the published minimum concentration required and 100x the concentration generally reported when DNP-INT is used to truncate PSI from the PETC (Khorobrykh and Ivanov 2002; Borisova-Mubarakshina, Naydov and Ivanov 2018; Krieger-Liszkay, Kienzler and Johnson 2000). Another important question was to assure the purity of the DNP-INT we had procured. Historically DNP-INT has only been available to researchers through collaboration with its creator, the late Achim Trebst. However, companies have recently started to synthesize and market this chemical as an inhibitor of PQ oxidation. Although our supplier, Cayman Chemicals, provided a full analytical analysis of their product to validate its purity, we purchased a second stock from a separate batch and decided to independently verify the structure of the product. The Turku University Instrumentation Center conducted ^1^H NMR on our DNP-INT stock, confirming the chemical’s structure and that the two benzene rings were connected (See Figure S2 Supplementary material).

To further test the reliability of our DNP-INT samples, and to confirm the proper functionality of the isolated thylakoid preparations, we decided to replicate published measurements that purportedly demonstrate effectiveness of DNP-INT in truncating PSI from the PETC (Khorobrykh and Ivanov 2002; Trebst and others 1978).

In this measurement, a Clark-type O_2_ electrode was used to compare Net O_2_ fluxes from isolated thylakoid samples (Figure 3), initially in the absence of artificial electron acceptors but with and without DNP-INT. The experiment was repeated with similar samples but MV was injected after one minute of illumination to observe its capacity to increase Net O_2_ uptake fluxes. The results (Figure 3.) faithfully reproduce the published outcome (Trebst and others 1978; Khorobrykh and Ivanov 2002). Compared to the untreated control (Black line), addition of DNP-INT (10 μM) to isolated thylakoid samples decreased rates of illuminated Net O_2_ uptake (Blue line). The addition of MV to the DNP-INT treated samples had no effect on this impaired flux (Pink line), in stark contrast to MV’s large effect of increasing the net O_2_ flux of untreated control samples (Red line). The fact that MV cannot increase O_2_ uptake rates in the presence of DNP-INT is traditionally interpreted as functional evidence for truncation of PSI from the PETC by 10 μM DNP-INT. That we could faithfully re-produce this result, with our stock of DNP-INT and our thylakoid material, in addition to independent confirmation via ^1^H NMR of the structure of our DNP-INT, we concluded that redox kinetics of PSI presented in Figure 2 were not the result of any artefacts and warranted in-depth investigation.

**Figure 3.**
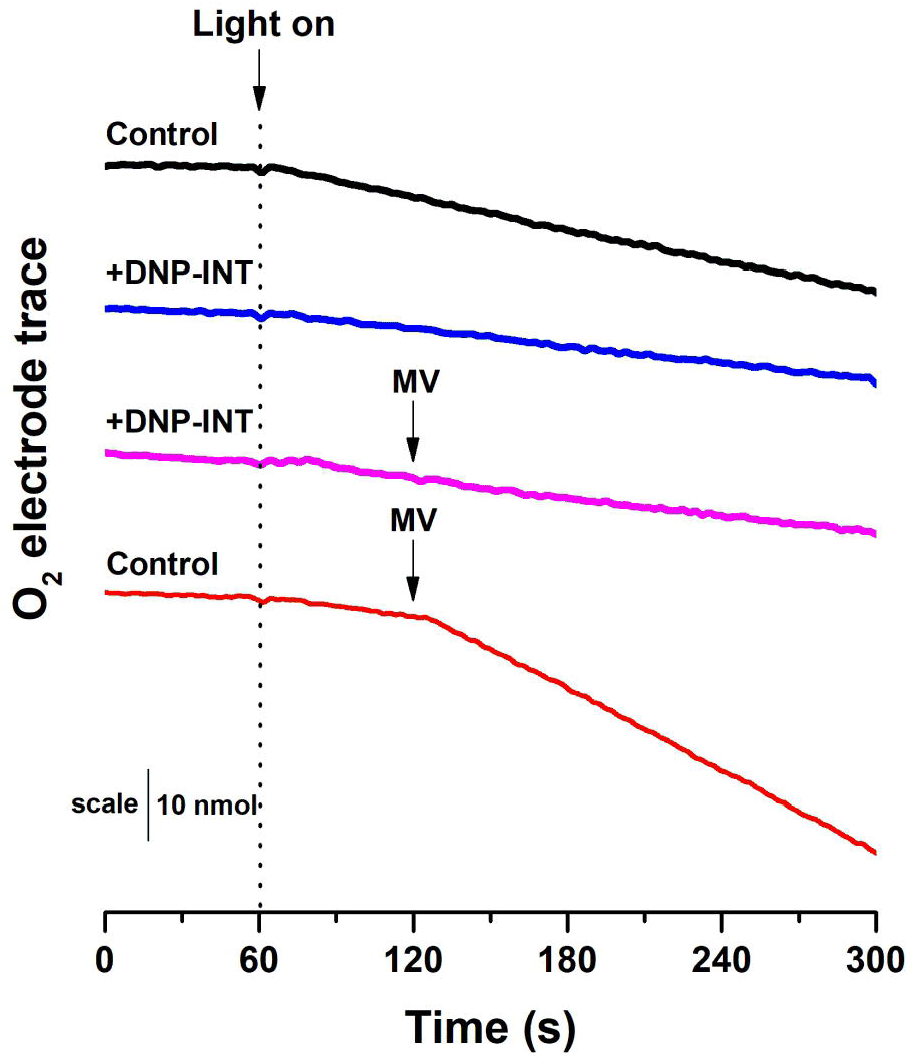
**Raw traces from O_2_ electrode** to demonstrate that our materials and methods can reproduce O_2_ electrode data historically used to show the efficacy in DNP-INT. Raw [O_2_] vs time traces were generated with thylakoids isolated from *Arabidopsis thaliana*. All measurements were conducted at a starting [O_2_] of approximately 250 nmol ml^−1^. Light intensity was 800 μmol photons m^−2^ s^−1^, Thylakoids equivalent to 15 μg Chl ml^−1^. Each curve is an average of n=6 replicate measurements.

### 3.4 MIMS data showed that O_2_ electrode assumptions are not always correct

Interpretation of data in Figure 2 surprised us by suggesting that DNP-INT did not fully inhibit the re-reduction of P700^+^ by actinic light pulses, as was expected and observed in samples incubated with DBMIB and DCMU (Figure 2). At the same time, the O_2_ electrode data in Figure 3 has been interpreted to imply that DNP-INT effectively blocks electron donation to MV (Figure 3) and therefore PSI is effectively truncated from the PETC by DNP-INT. We inferred the best explanation for this discrepancy to rest with the inherent limitations of the Clark-type O_2_ electrode, which can only measure Net O_2_ fluxes. This is particularly problematic when the stoichiometry of O_2_ production and consumption is not clearly known, as is the case when O_2_ is both produced and consumed in the same sample, at unknown relative rates depending on the characteristics of the O_2_ reduction pathways active under different conditions (see Figure 1). To investigate whether this limitation of O_2_ electrodes has potentially contributed to miscalculate/misinterpret the efficacy of DNP-INT’s activity in isolated thylakoid samples, we employed the resolving power of MIMS. MIMS is able to discriminate between O_2_ isotopologues to independently and simultaneously measure activity of PSII, and concomitant reduction of O_2_ in isolated thylakoids (Furbank and Badger 1983). By independently measuring the activity of PSII it was possible to unambiguously judge whether MV added to the DNP-INT inhibited samples could increase the rate of O_2_ evolution or not, providing a clear test of whether or not PSI is truncated from the PETC in isolated thylakoids. As a final test of the capacity for DNP-INT to impair electron donation to PSI in intact leaves, we used the MIMS to measure whether or not leaf discs infiltrated with DNP-INT could still fix CO_2_ in comparison to those infiltrated with DBMIB.

In both isolated thylakoid and leaf disc samples, PSII activity was measured as the Gross O_2_ evolution rate, (or production of ^16^O_2_ derived from PSII splitting of H_2_^16^O (see Figure 1)). The reduction of O_2_ was measured directly through enrichment of the samples with the stable ^18^O_2_ isotope, which following an N_2_ purge to reduce ^16^O_2_ became the primary electron acceptor in the isolated thylakoid samples. Leaf disc samples were purged with N_2_ gas before being enriched to approximately 2% with ^13^CO_2_, used as the terminal electron acceptor, and 3% ^18^O_2_ to monitor O_2_ reduction rates. Photorespiration in leaf discs was minimized by maintaining an elevated CO_2_ partial pressure, which left mitochondrial respiration and potentially the Mehler reaction as primary O_2_ uptake pathways. Monitoring ^12^CO_2_ efflux rates derived from mitochondrial respiration enabled an estimate of illuminated respiration rates, minus any CO_2_ re-fixation (Busch 2018) to offset its contribution to ^18^O_2_ consumption, whilst the latter was minimized through low O_2_ partial pressure.

MIMS data from isolated thylakoid samples are presented in Figure 4A-D. When illuminated at 120 μmol photons m^−2^ s^−1^ little difference in PSII activity (Blue lines, Figure 4) could be observed between control and DNP-INT treated thylakoids (Figure 4A,C). However, Gross O_2_ uptake rates (Red Lines, Figure 4) were slightly lower in the latter, resulting in Net O_2_ fluxes (Grey Lines, Figure 4), calculated from the difference between the two Gross O_2_ rates, being smaller (less negative) in the DNP-INT samples. The difference in Net O_2_ fluxes matches the results from the Clark-type O_2_ electrode in Figure 3. However, the observation that O_2_ evolution rates were actually very similar between the two conditions was unexpected. The smaller O_2_ uptake rates in DNP-INT treated samples (Figure. 4C) suggests that less H_2_O_2_ accumulated in the presence of this chemical.

**Figure 4.**
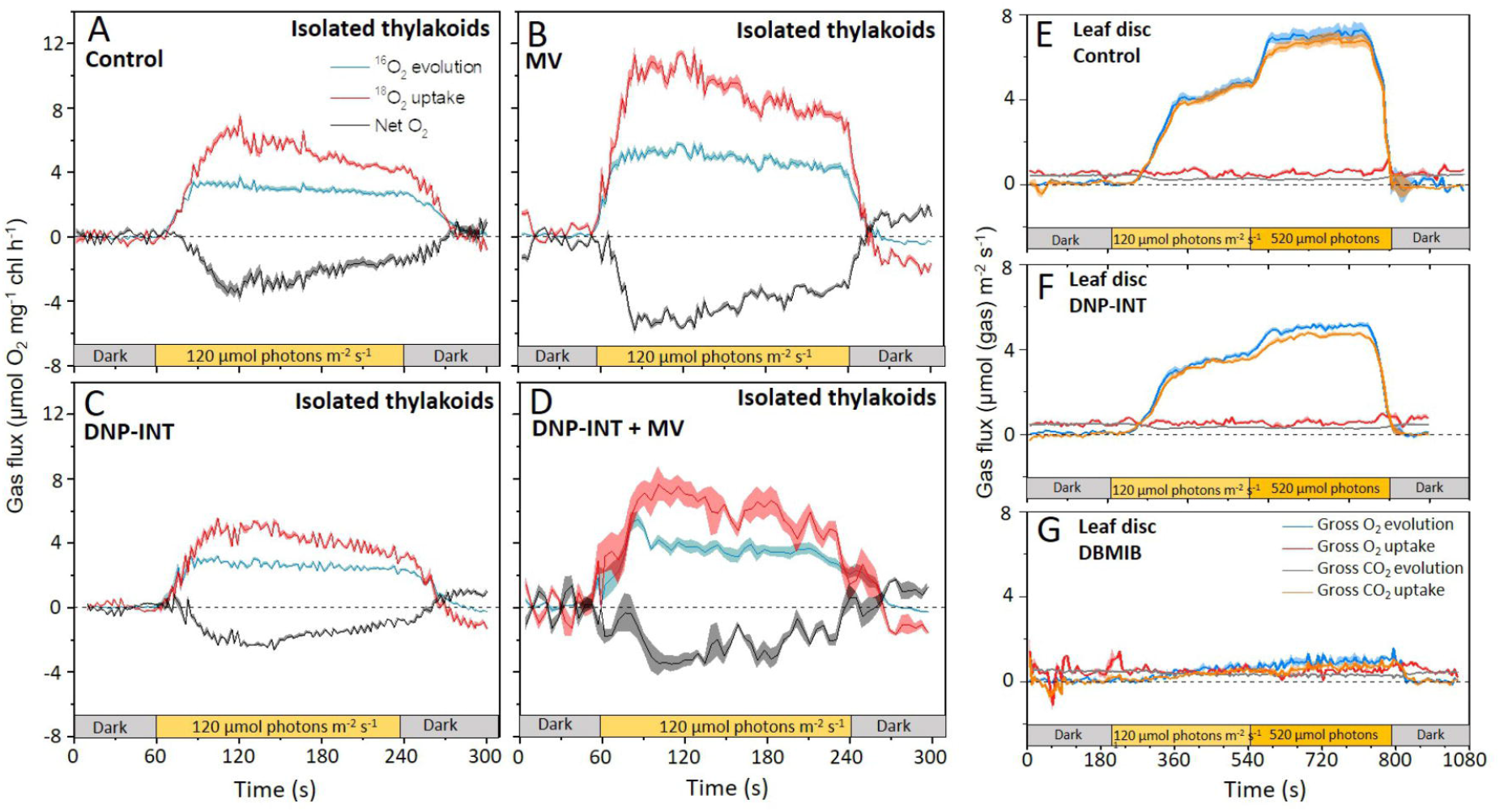
Integrated gas flux versus time plots of isolated thylakoid and intact leaf disc samples (±SE) Simultaneously measured rates of O_2_ Evolution and Consumption from isolated thylakoids (50 μg chlorophyll ml^−1^) **(A-D)**, with the addition of CO_2_ consumption and production rates from intact 12.5 mm leaf discs **(E-F)** measured with MIMS. Thylakoid samples measured under similar conditions used in Dual-PAM, see Figure 2. Leaf discs were floated overnight in water, or water containing 10 μM DNP-INT or 10 μM DBMIB in darkness. The rates of all isolated thylakoids were offset to zero in pre-illuminated darkness. Isolated thylakoid curves comprise averaged data from n=7 (Control), n=8 (DNP-INT), n=8 (MV) and n=4 (DNP-INT+MV) independent replicates. Leaf disc curves comprise the averaged data of n=4 (control), n=4 (DNP-INT) n=3 (DBMIB) independent replicates.

Addition of MV to thylakoids (Figure 4B) resulted in a large increase to rates of PSII O_2_ evolution and Gross O_2_ uptake, compared to the control, demonstrating that PSII activity is dictated by the strength of the acceptors in isolated thylakoid samples. The proportionate increase in both O_2_ fluxes resulted in an increased difference between them, producing a larger negative Net O_2_ flux, which also strongly reflected the Clark-type O_2_ electrode data in Figure 3. Whilst it was clear that DNP-INT impeded rates of MV photoreduction (Figure 4D), it was also clear that MV increased rates of PSII activity in the DNP-INT inhibited samples (for a direct comparison refer to supplementary Figure S3 where we plot the Gross O_2_ evolution rates measured under the four conditions on the same axis). This suggests that in-spite of the presence of DNP-INT, MV was able to increase the acceptor side capacity, which strongly supports the conclusions from P700 data in Figure 2, implying that DNP-INT does not truncate PSI from the PETC. In addition, the data shows that O_2_ uptake rates did not increase proportionately with the increased PSII activity. This resulted in a smaller increase in the negative Net O_2_ flux than would be predicted from a measure of PSII O_2_ evolution alone, which again fits the O_2_ electrode data in Figure 3 and suggests that DNP-INT treated samples seem to generate less stable H_2_O_2_ than samples lacking this compound. Overall, deconvolution of O_2_ generation and uptake afforded by MIMS has revealed that the Net O_2_ fluxes observed do not necessarily correlate to the activity of PSII for a given condition. This suggests that the Net rates reported by Clark-type O_2_ electrodes are unable to provide sufficient information to accurately interpret the site-specific complexities of ROS formation and scavenging in isolated thylakoid membrane samples lacking artificial donors and acceptors.

The final experiment to test the efficacy of DNP-INT to block electron transport to PSI in infiltrated leaves was to measure the capacity of intact leaves, infiltrated with 10 μM DNP-INT, to fix CO_2_ during illumination using MIMS. Again, DBMIB was used as a comparison when electron transport at the Cyt-*b6f* complex was significantly inhibited. After five minutes darkness, leaf discs were illuminated five minutes at 120 μmol photons m^−2^ s^−1^, then five minutes at 520 μmol photons m^−2^ s^−1^ before extinguishing the actinic light. As expected, photosynthetic activity was severely impaired by infiltration with DBMIB (Figure 4G) compared to untreated controls (incubated in water, Figure 4E). However, samples incubated with DNP-INT (Figure 4F) exhibited only a small reduction, of approximately 30%, in CO_2_ fixation rates during illumination at both irradiances. In all samples, rates of mitochondrial respiration (CO_2_ efflux, grey lines Figure. 4E-G) were similar.

## 4. Conclusions

In efforts to assess the contribution of PSI-independent O_2_ reduction pathways operating during illumination within the PETC, it is necessary to disconnect PSI completely from the rest of the PETC with effective inhibitors. Whilst DCMU and DBMIB functioned as expected in this regard, we were surprised to find that DNP-INT failed to block re-reduction of P700^+^ at concentrations up to 1000x the published minimum concentration required. We confirmed the structure of our DNP-INT stocks through ^1^H NMR, and we were able to reproduce a published O_2_ electrode experiment used historically to support the contention that DNP-INT effectively truncates PSI from the PETC in isolated thylakoid samples. However, we observed with a Dual-PAM that when compared to the inhibitors DCMU and DBMIB, DNP-INT was unable to decrease, but not block actinic reduction of P700^+^. This suggested that DNP-INT can only slow electron transfer through the PETC in isolated thylakoid samples, enabling the Mehler reaction, and in intact system, the carbon reduction cycle, to continue functioning. Using MIMS to independently and simultaneously measure the rates of O_2_ production and consumption in isolated thylakoids, we demonstrated that MV added to DNP-INT treated samples was able to increase rates of PSII activity. However, the O_2_ uptake rate did not increase commensurately, potentially due to unknown complexities of ROS scavenging within thylakoid membranes. As such, the Net O_2_ flux did not appear to be different from the DNP-INT treated samples lacking MV. Through these experiments, we demonstrated that measurements of Net O_2_ fluxes with an O_2_ electrode are insufficient for measurements of the site-specific O_2_ reduction pathways operating within thylakoid membranes, due to the complexities of the competing routes of superoxide formation and quenching that potentially invalidate some assumptions underpinning the interpretation of Net O_2_ fluxes. Finally, we showed that leaf discs incubated with DNP-INT could still fix CO_2_, at rates impaired by approximately 30%, whereas DBMIB severely impaired all photosynthetic fluxes from similar samples. We suggest that DNP-INT should no longer be used as an inhibitor of PSI reduction, and that results reliant on the assumption that DNP-INT had effectively inhibited the activity of PSI must be revisited. Nonetheless, the chemical may still find use for its apparent ability to slow electron transport between the PQ pool and PSI. This could be useful as a tool to artificially replicate the induction of photosynthetic control, imparted by the Cyt-*b*_*6*_*f* complex by an acidified lumen, potentially in the absence of NPQ or state transitions. It may also find use with researchers trying to modulate electron transfer rates in biophotovoltaic applications (Tschörtner, Lai and Krömer 2019). Finally, the disagreement in results between characterization studies of DNP-INT in purified complexes and our findings from intact samples may point to currently unknown aspects of cyt-b6f function. Perhaps DNP-INT only partially blocks cyt-b6f due to competitive inhibition with some other substrate that may now be studied with DNP-INT. Or perhaps the results reflect a sub-population of DNP-INT sensitive cyt-b6f which may correlate to some spatial distribution in the thylakoid membrane, or its participation in super or mega complexes. Further work to explain the apparent discrepancies of DNP-INT function between spectroscopic measurements in highly purified samples and our results obtained with isolated thylakoids and intact leaf discs may yield significant information about photosynthesis in the future.

## Author contribution

AT, DF and EMA conceptualized and designed the work. DF and AT carried out the experimental work, data analysis and interpretation of data. DF drafted the manuscript. AT and EMA revised the content for final submission.

## Acknowledgements

Thanks to Tuomas Karskela at the Turku instrument Centre for ^1^H NMR measurement and analysis.

