## Supplementary figures and images for "A commonly used photosynthetic inhibitor fails to block electron flow to photosystem I in intact systems"

### Figure S1

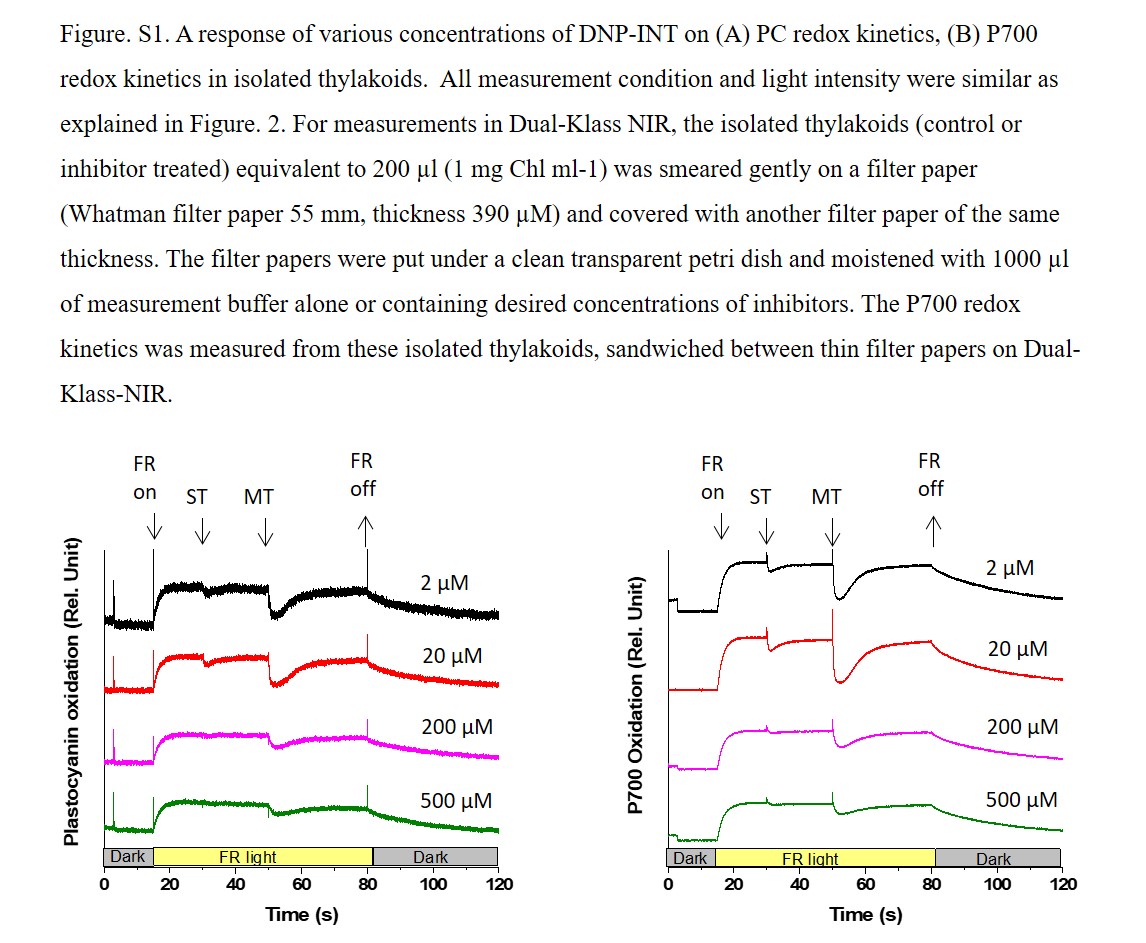

### Figure S2

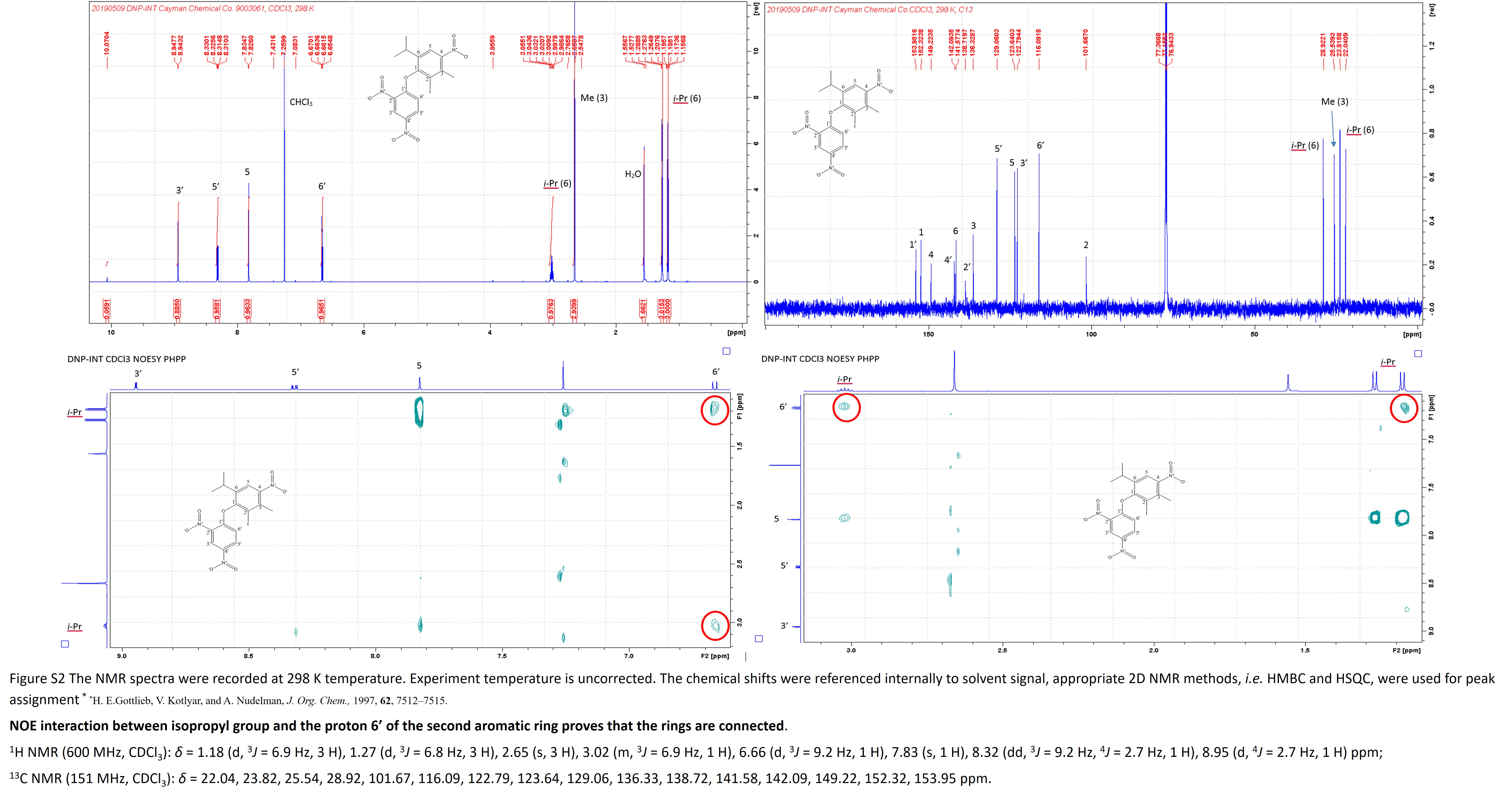

### Figure S3

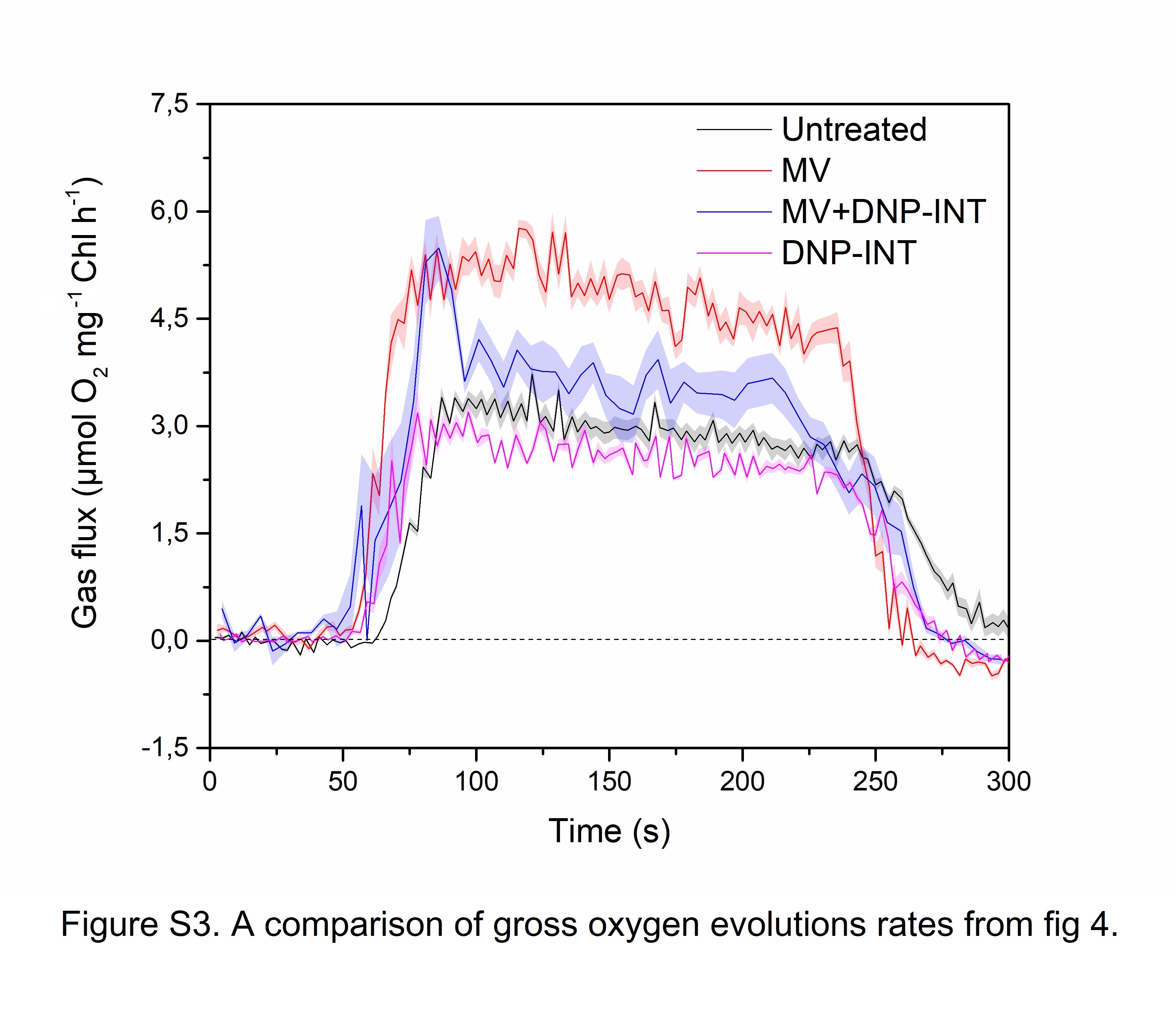
